# Peptide Ancestry Informative Markers in Uterine Neoplasms from Women of European, African and Asian Ancestry

**DOI:** 10.1101/2020.11.23.380337

**Authors:** Nicholas W. Bateman, Christopher M. Tarney, Tamara S. Abulez, Brian L. Hood, Kelly A. Conrads, Ming Zhou, Anthony R. Soltis, Pang-ing Teng, Amanda Jackson, Chunqiao Tian, Clifton L. Dalgard, Matthew D. Wilkerson, Michael D. Kessler, Zachary Goecker, Jeremy Loffredo, Craig D Shriver, Hai Hu, Michele Cote, Glendon J. Parker, James Segars, Ayman Al-Hendy, John R. Risinger, Kathleen M. Darcy, Yovanni Casablanca, G. Larry Maxwell, Thomas P. Conrads, Timothy D. O’Connor

## Abstract

Characterization of ancestry-linked peptide variants in disease-relevant patient tissues represents a foundational step to connect patient ancestry with molecular disease pathogenesis. Nonsynonymous single nucleotide polymorphisms (SNPs) encoding missense substitutions within tryptic peptides exhibiting high allele frequencies in European, African, and East Asian populations, termed peptide ancestry informative markers (pAIMs), were prioritized from 1000 genomes. *In silico* analysis shows that as few as 20 pAIMs can determine ancestry proportions similarly to >260K SNPs (R^2^=0.9905). Multiplexed proteomic analysis of >100 human endometrial cancer cell lines and uterine leiomyoma tissues resulted in the quantitation of 62 pAIMs that correlate with self-described race and genotype-confirmed patient ancestry. Candidates include a D451E substitution in GC vitamin D-binding protein previously associated with altered vitamin D levels in African and European populations. These efforts describe a generalized set of markers for proteoancestry assessment that will further support studies investigating the impact of ancestry on the human proteome and how this relates to the pathogenesis of uterine neoplasms.

## Introduction

Proteogenomics aims to integrate protein-level measurements with companion transcriptome and genome sequencing data (1) to elucidate complex systems-level relationships between protein and transcript-level expression including identification of proteins encoding missense substitutions that may impact protein function (1, 2). Understanding the impact of proteogenomic alterations on disease risk and patient outcome is a priority for population-level investigations (3), particularly those focusing on cancer, such as efforts by The Cancer Genome Atlas (TCGA) (4) and the recently initiated Applied Proteogenomics OrganizationaL Learning and Outcomes (APOLLO) research network (5). Many historic efforts have poor representation of U.S. minorities amongst patient cohorts analyzed (6) and there remains a paucity of systems-level molecular analyses within powered cohorts to elucidate race-related differences in disease etiology. In the context of gynecologic disease, patients with uterine leiomyomas (fibroids) and endometrial cancers experience significant racial disparities in disease incidence and outcome (7–13). Among possible molecular mechanisms that may give rise to racial disparities in gynecologic disease, one such manifestation could be from non-synonymous single nucleotide polymorphisms (nsSNPs) encoding missense substitutions that may alter protein function(s). Defining ancestry-linked peptide variants in disease-relevant cells and tissues will augment proteogenomic discovery efforts that aim to define molecular drivers underlying racial disparities in gynecologic diseases.

Ancestry informative markers (AIMs) comprise SNPs that are highly differentiated within discrete ancestral populations (14–16). Most of these markers occur within non-protein coding regions, but with large-scale sequencing now available for a diversity of ancestries (17, 18), especially in protein coding regions (19, 20), it is now possible to assess AIMs that are detectable through proteomics data alone. To this end, efforts to date have identified genetically variant peptides differentiating individuals of European and African descent (21) as well as Asian populations (22) and these have been explored for forensic human identification, such as through proteomic analyses of human hair and bone samples. We leveraged population-level genotyping data from the 1000 Genomes Project (23) to identify SNPs encoding missense substitutions that exhibit altered allele frequencies with European, African, and East Asian ancestries and prioritized candidates occurring within tryptic peptides observable by routine, bottom-up proteomic workflows to explore these within disease-relevant models and tissues from uterine neoplasms. Although these candidates are not tissue dependent and represent a generalized set of markers for proteoancestry assessment in diverse tissues and cell lines, we show here the ability to ancestrally characterize endometrial cancer cell line models as well as uterine fibroid tissues from women representing diverse racial and ancestral backgrounds using multiplexed proteomic approaches. Our study further defines a foundational set of ancestry-linked variant peptides within uterine tissues that will support ongoing efforts to investigate germline as well as somatic proteogenomic alterations underlying ancestry-linked disease biology and how this may further relate to racial disparities in the pathogenesis of uterine neoplasms.

## Results

### Selection and *In Silico* Analyses of Peptide Ancestry Informative Markers (pAIMs)

We selected >640K SNPs from the 1000 Genomes Project (23) encoding nonsynonymous substitutions and further filtered them to those exhibiting ≥ 50% allele frequency differences between individuals of European (i.e. FIN, GBR, IBS, and TSI; N=404), African (i.e. ESN, GWD, LWK, MSL, and YRI; N=504), and East Asian (i.e. CDX, CHB, CHS, and KHV; N=400) descent. Candidates were also filtered to prioritize missense substitutions occurring in tryptic peptides (≥6 and ≤40 amino acids in length) that would be observable within “bottom-up”, multiplexed analyses of disease-relevant cell lines and tissues collected from individuals representing diverse ancestries (Workflow Figure 1A, Supplemental Table 1A). This filtering approach resulted in 1,037 peptide ancestry informative markers (pAIM) mapping to 831 unique proteins spanning all 22 autosomes and exhibit the greatest frequencies on Chr1 (12.3%) and Chr19 (7.5%) (Supplemental Table 1A). Most of these parent proteins encode a single pAIM (Supplemental Table 1A) with a subset of 136 candidates encoding ≥2 pAIMs as well as 3 candidates encoding >5 pAIMs; filaggrin (10 pAIMs), cardiomyopathy-associated protein 5 (7 pAIMs) and HLA class II histocompatibility antigen, DP alpha 1 chain precursor (6 pAIMs). Functional enrichment analyses of proteins encoding pAIMs revealed significant enrichment of cellular pathways regulating the cellular matrisome as well as keratinocyte differentiation and sensory perception (Supplemental Table 1B). Unique pAIMs exhibiting ≥50% allele frequencies were most prevalent in African populations (344 total sites) followed by East Asian (229) and European (79) ancestries and East Asian as well as European populations exhibited nearly 2-fold greater conservation (208) of shared pAIMs relative to African populations (92 and 85 respectively) (Figure 1B). Fixation index (Fst) values for nonsynonymous single nucleotide polymorphisms (SNPs) encoding pAIMs exhibited an average of 0.312 ± 0.1 consistent with Fst threshold expectations for identity-informative SNPs used in forensic DNA sequencing (24) (Supplemental Table 1A). The clinical significance and known disease pathogenicity of SNPs encoding pAIMs within the ClinVar resource was assessed for 1,031 subset pAIMs mapping to the Ensemble Variant Effect Predictor tool (25). Most of these pAIMs had no previous associations with clinical significance. We did however identify 87 pAIMs of unknown clinical significance that map to putative functional protein domains (Figure 2A, Supplemental Table 1A). The remaining candidates were largely undocumented for disease association in ClinVar with seven pAIMs candidates correlating with altered drug responses or as being risk factors for disease. This latter subset included a substitution in kinesin family member 6 that is associated with altered response to statin treatment (26) (W719R, rs20455) and has higher allele frequencies within African (86%) and East Asian (53%) populations, or as being risk factors for several diseases (Supplemental Table 1A). Risk factors included a substitution in aurora kinase A (F31I, rs2273535) which has higher allele frequencies within East Asians (67%) and is associated with an increased risk of developing colon cancer within these populations (27). We also assessed the impact of pAIM substitutions on protein function via *in silico* prediction analyses using SIFT (28), Poly-Phen-2 (29), and PROVEAN (30) and identified 23 pAIMs that may be deleterious (Figure 2B). Although many of these candidates have unknown clinical significance, five candidates have been shown to not be pathogenic for disease, i.e. categorized by ClinVar as benign or likely benign (Figure 2C, Supplemental Table 1A). These candidates include a substitution in ATP-binding cassette transporter sub-family C member 11 (G180R, rs17822931) that has high allele frequencies within East Asian populations (75%) and is significantly correlated with Axillary Osmidrosis in Chinese Han populations (31) as well as increased risk for breast cancer in Japanese women (32). Further analyses of these 23 putatively deleterious pAIMs revealed they exhibit higher allele frequencies in East Asian versus European and African populations (Figure 2D). We also utilized as a test set of populations CEU, a European population from Utah; ASW, an admixed African-American population from the South West; ACB, an admixed African Caribbean population; and JPT, a Japanese population from East Asia to assess the accuracy of pAIMs to classify ancestral proportions relative to standard genotype estimates *in silico*. We apply a random sampling analyses of nsSNPs encoding pAIMs and compared them to using 266,403 LD-pruned SNPs by standard estimates approaches. We find that as few as 20 variants can recapitulate genome-wide ancestry proportions, using standard approaches (33–35) (error is increased due to smaller feature set, but average of 500 random sampling assessments remains robust is R^2^>0.99) (Figure 3).

**Figure 1:**
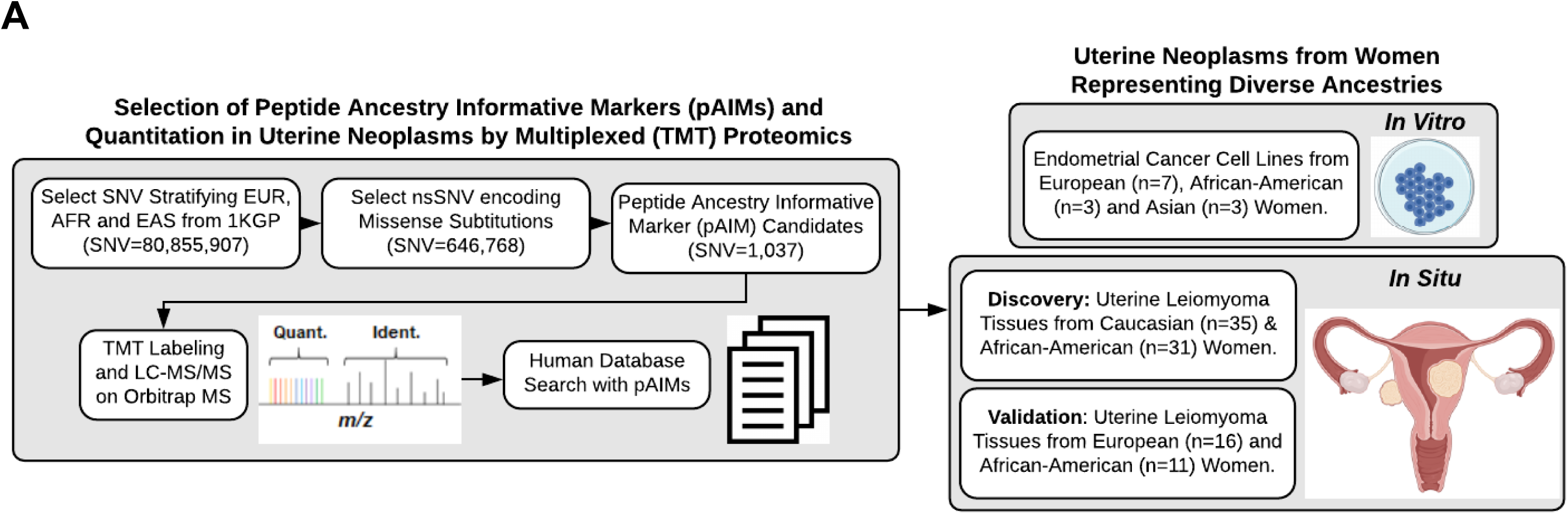
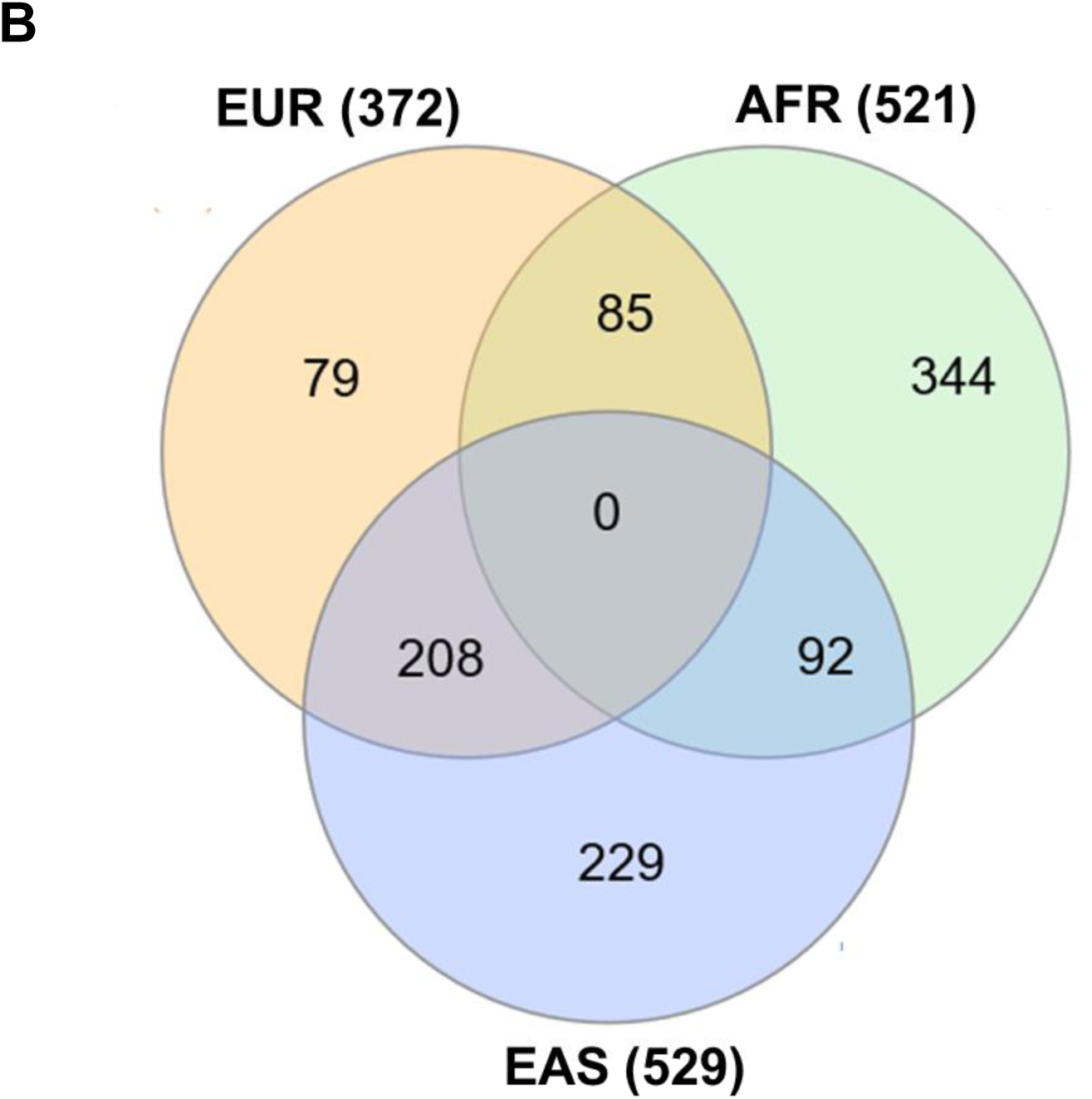
Selection of Peptide Ancestry Informative Markers (pAIMs) and Workflow for Analyses in Cell Lines Models and Tissues of Uterine Neoplasms collected from Women of Diverse Ancestries. A: >640K single nucleotide polymorphisms (SNPs) were filtered to prioritize 1,037 missense substitutions occurring within lysine or arginine-terminating tryptic peptides, ≥ 6 and ≤ 40 amino acids in length, “so-called” peptide ancestry informative markers (pAIMs). We then investigated pAIMs in endometrial cancer cell line models as well as uterine leiomyoma tissues from women representing diverse ancestries. B: Comparison of pAIMs exhibiting ≥ 50% allele frequencies within individuals of European (EUR), African (AFR) or East Asian (EAS).

**Figure 2:**
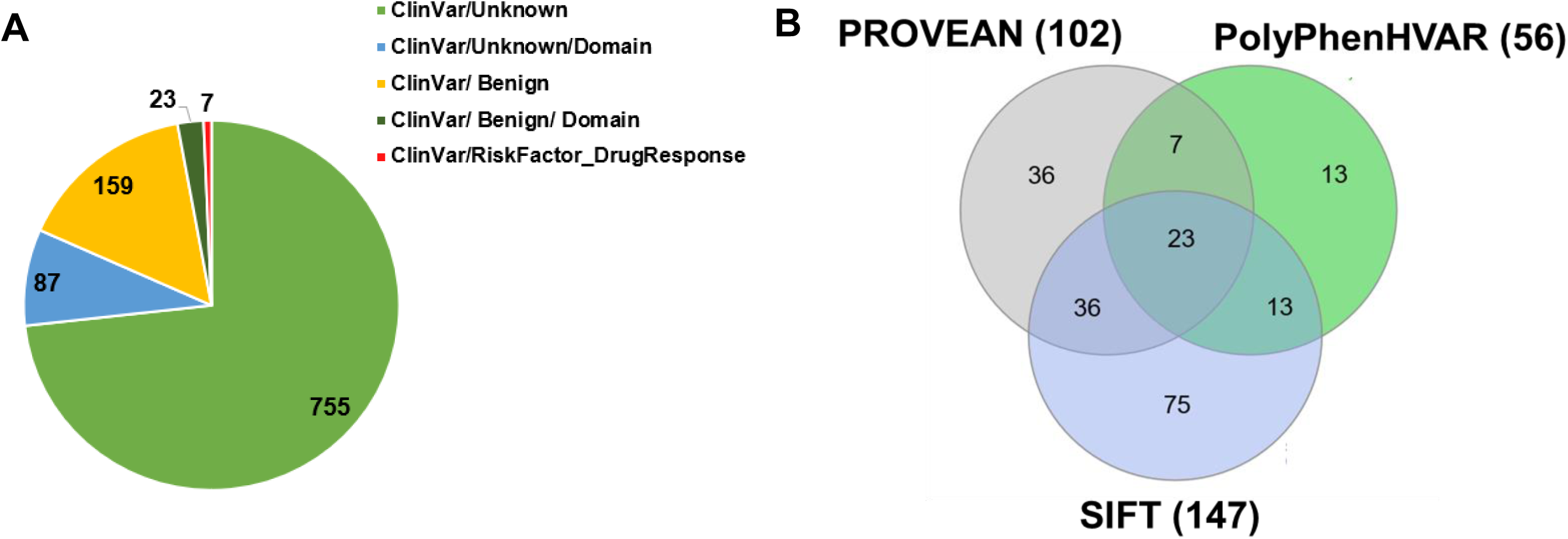
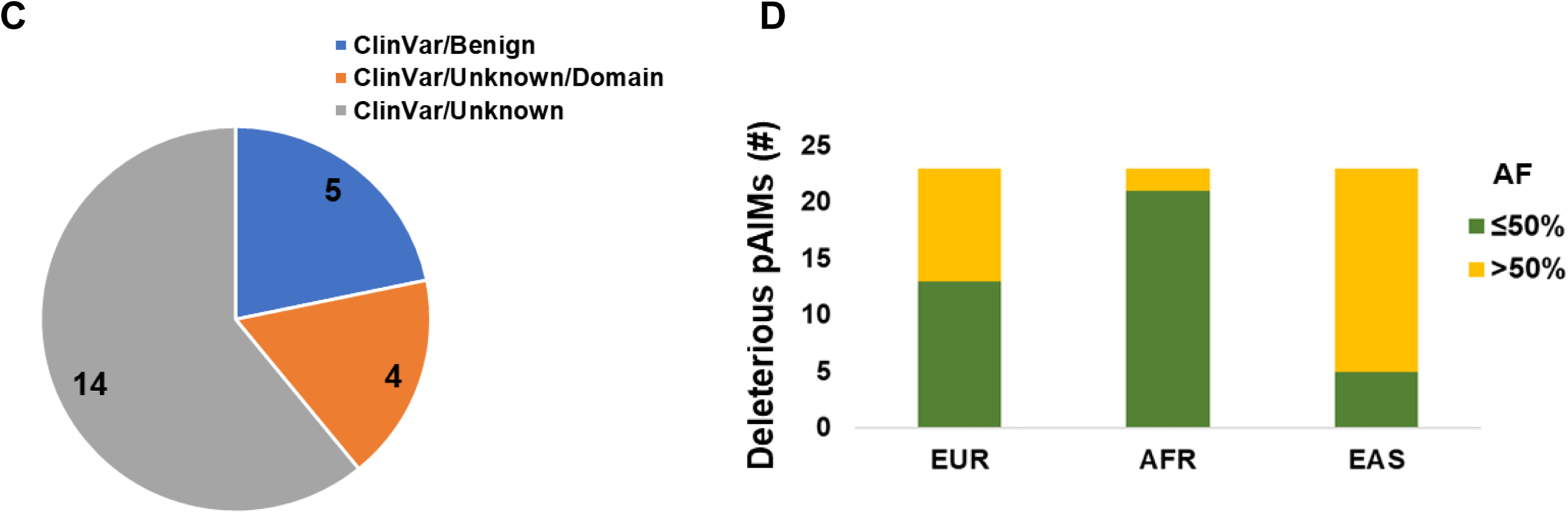
*In silico* analyses of Peptide Ancestry Informative Markers (pAIMs) to assess Clinical Significance, Localization to Protein Domains as well as to Predict Impact on Protein Function. A: Clinical significance (ClinVar) and protein domain (Uniprot) localization of 1,037 pAIMs. B: Comparison of pAIMs predicted to be deleterious to protein function by PROVEAN, PolyPhen and SIFT functional prediction tools. C: Clinical significance (ClinVar) and protein domain (Uniprot) localization of pAIMs predicted to be deleterious to protein function. D: Allele frequency of pAIMs predicted to be deleterious to protein function in European (EUR), African (AFR) or East Asian (EAS) populations

**Figure 3:**
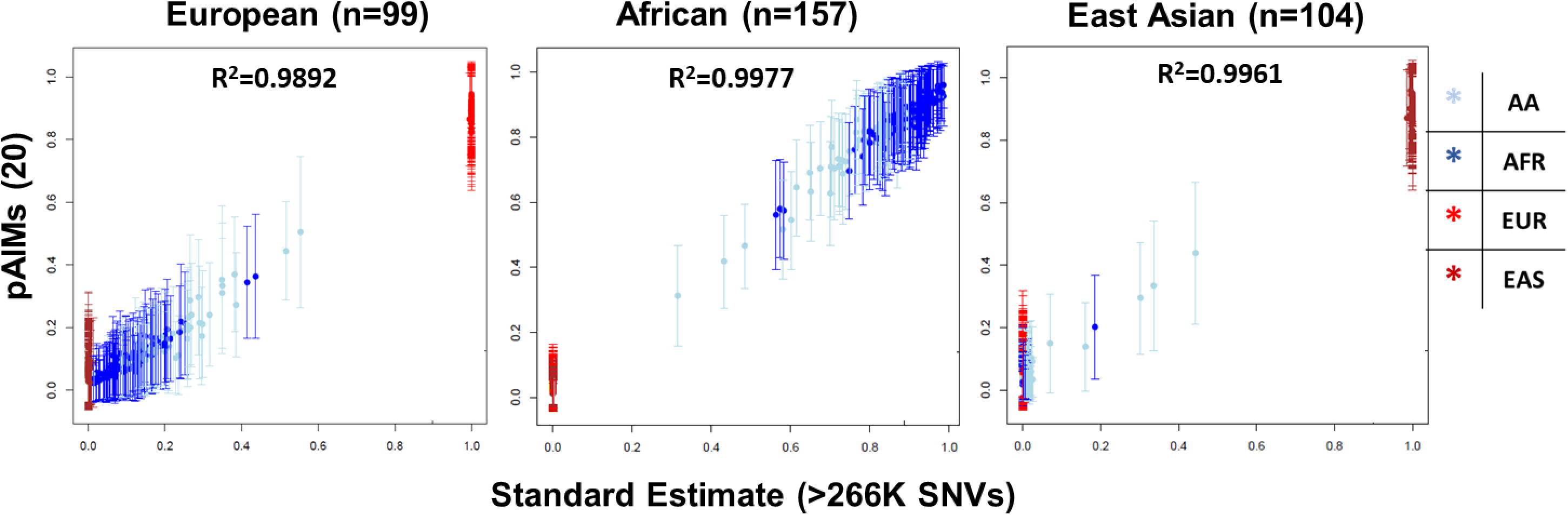
Comparison of European (EUR), African (AFR) or East Asian (EAS) Global Ancestry Classification using Peptide Ancestry Informative Markers (pAIMs) with Standard Estimates. Random sampling of pAIMs in European, African and East Asian reference populations from 1K genome project revealed that a minimum of 20 pAIM-encoding nonsynonymous single nucleotide polymorphisms (SNPs) can classify global ancestry with accuracy comparable to standard estimates using >266,000 SNPs, i.e. R^2^=0.9943 ± 0.005.

### Quantitation of Peptide Ancestry Informative Markers (pAIMS) in Endometrial Cancer Cell Lines

We determined the ancestry of thirteen endometrial cancer cell line models by standard estimates using comparison of whole genome sequencing (WGS) data to reference populations (Table 1). Most EC models were established from women of European descent (n=7), with a subset corresponding to women of African (n=3) and East Asian (n=3) descent. We performed quantitative bottom-up proteomic analyses employing a multiplexed tandem-mass tag (TMT) approach and quantified 133,473 total peptides that included 43 high-confidence pAIM variant peptides (Supplemental Table 2), where MS2 ions flanking the pAIM substitution of interest were confirmed using SpectrumAI (36). Among these pAIMS, 39 were quantified in >50% of all EC cell lines and this feature subset was assessed to investigate pAIM abundance relative to cell line ancestries and genotypes for the nsSNP of interest. Of these pAIM candidates, 30 are of unknown clinical significance, 12 are undocumented for disease association in ClinVar and one candidate (rs8010699) is of uncertain significance for disease. This latter variant encodes a substitution in nesprin-2 (H3309R) that is associated with Emery-Dreifuss muscular dystrophy and has further been correlated with expression of the tumor suppressor cyclin-dependent kinase inhibitor 1 (CDK1/ p21) in HBV-related hepatocellular carcinomas (37). A subset of 17 pAIMs exhibited >50% allele frequency within European ancestry populations and we found that these candidates were significantly elevated in cell lines from individuals of European versus African ancestry (Mann Whitney U (MWU) Median = +1.003 median fold difference, p=0.0333) as well as East Asian ancestry (East Asian ancestry Median = +0.57 median fold difference, p=0.0167) (Figure 4A). We further find that these European ancestry-correlated pAIMs were most abundant in European ancestry cell lines that were homozygous and heterozygous for the pAIM variant allele (+/+ and -/+ genotype) relative to those homozygous for the reference allele (-/-) (MWU p=0.0006). Another subset of 21 pAIMs quantified exhibited >50% allele frequency within African ancestry populations and were significantly elevated in cell lines from individuals of African versus European (MWU p=0.0333) ancestry and trended as elevated in African versus East Asian ancestry cell lines (+1.0676 median fold difference) (Figure 4B). These African ancestry-associated pAIMs were most abundant in African cell lines homozygous and heterozygous for the pAIM variant allele (+/+ and -/+ genotype) relative to those homozygous for the reference allele (-/-), i.e. +/+ vs. -/- cell lines = +1.69 median fold difference. We also quantified 17 pAIMs exhibiting >50% allele frequency within East Asian populations and found that these are elevated in cell lines from individuals of East Asian versus European ancestry (+0.4058 median fold difference) and African (+0.84563 median fold difference) ancestry (Figure 4C). These East Asian ancestry-related pAIMs further tracked with genotype, i.e. (+/+ pAIM allele versus -/- cell lines = +1.99 median fold difference).

**Figure 4:**
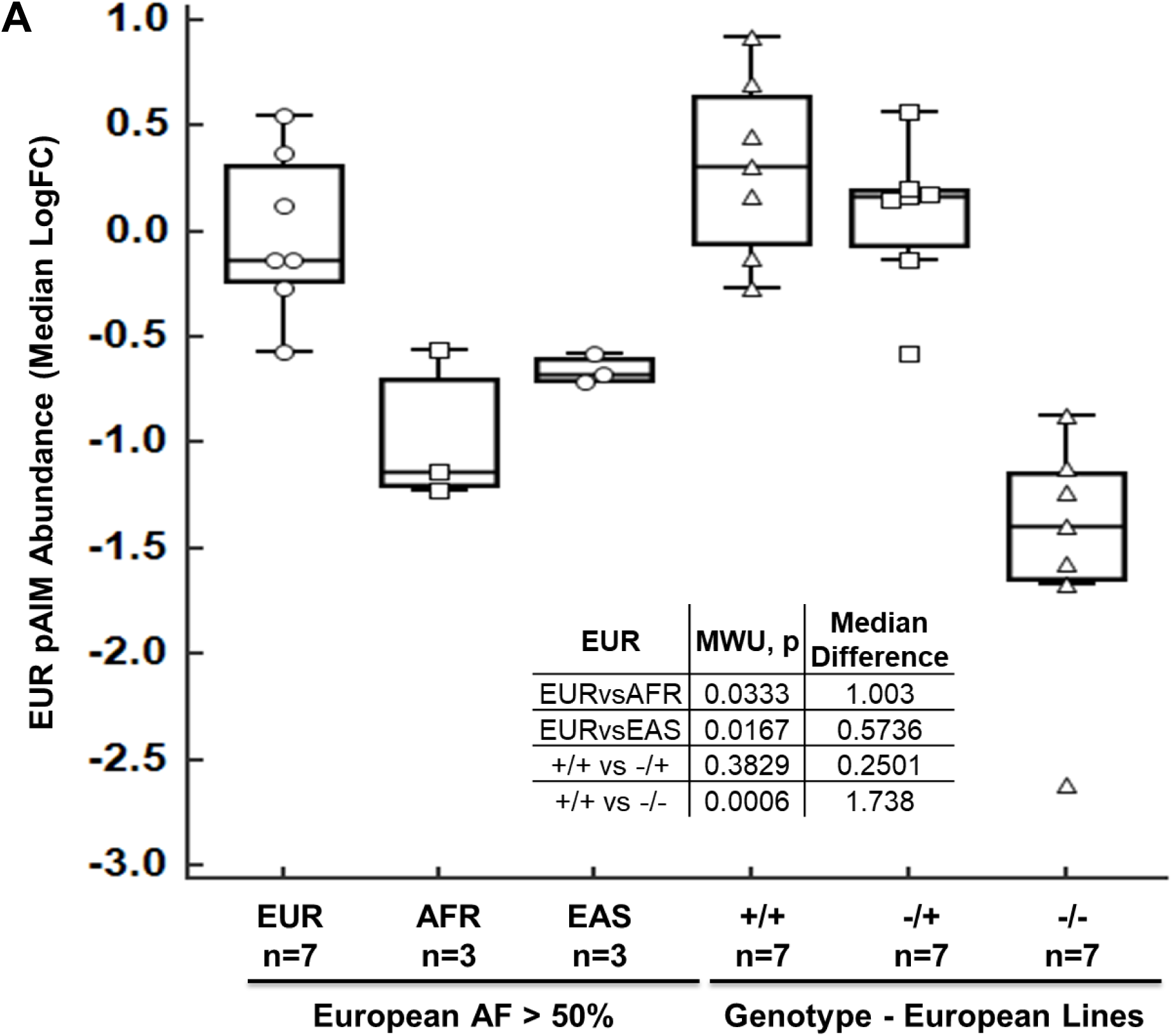
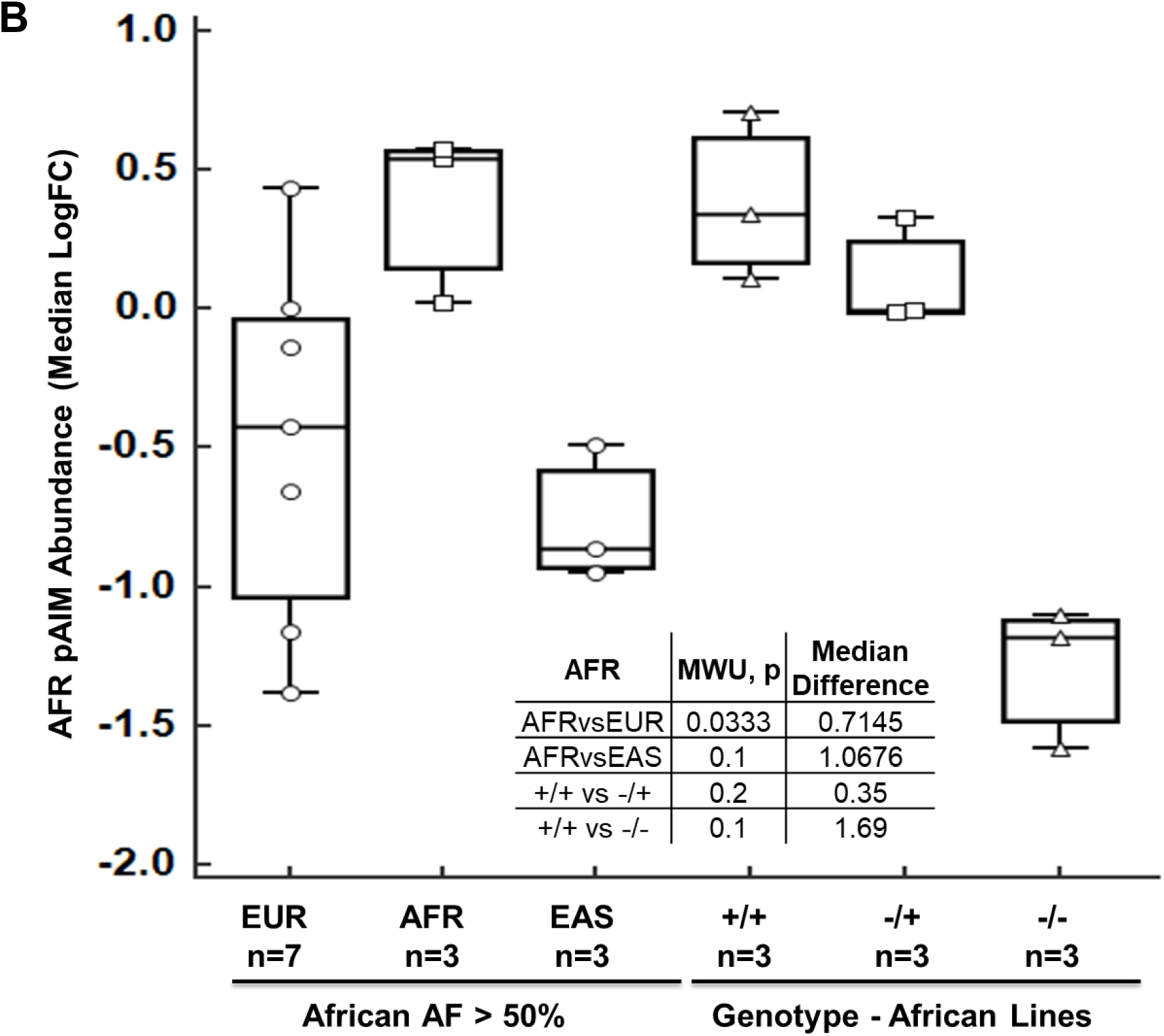
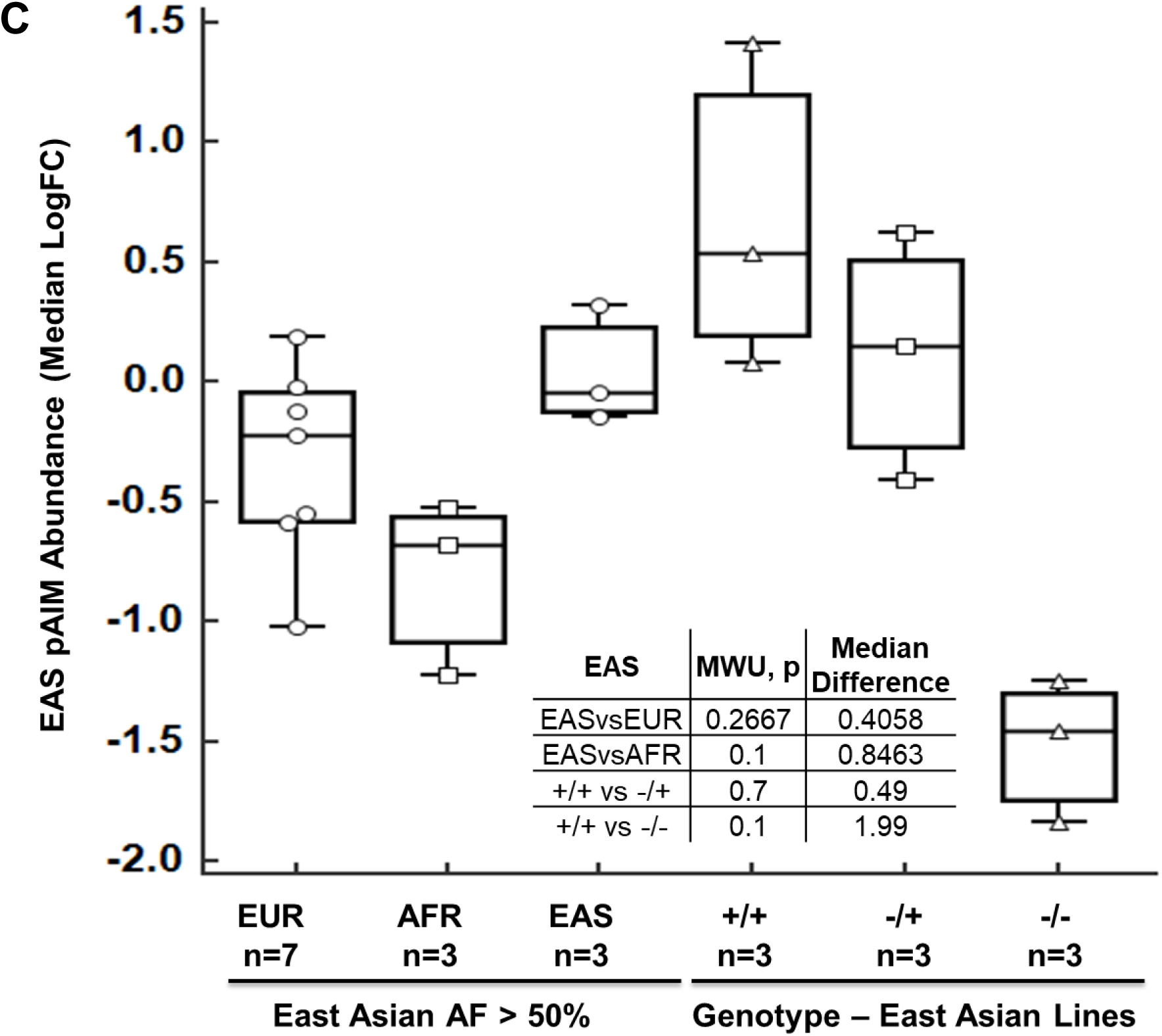
Quantitation of Peptide Ancestry Informative Markers (pAIMs) within Endometrial cancer cell Lines. A: Box plots detail the median logFC abundance of 17 EUR pAIMs within cell lines stratified by European (EUR), African (AFR) or East Asian (EAS) ancestry as well as by pAIM variant allele genotype observed by companion whole genome sequence analyses of EUR cell lines. B: Box plots details the median logFC abundance of 21 AFR pAIMs within cell lines stratified by European (EUR), African (AFR) or East Asian (EAS) ancestry as well as by as well as by pAIM variant allele genotype observed by companion whole genome sequence analyses of AFR cell lines. C: Box plots detail the median logFC abundance of 17 EAS pAIMs within cell lines stratified by European (EUR), African (AFR) or East Asian (EAS) ancestry as well as by pAIM variant allele genotype observed by companion whole genome sequence analyses of EAS cell lines.

**Table 1:** Endometrial Cancer Cell Lines Analyzed by Whole Genome Sequence and Quantitative Proteomic Analyses. Cell lines include both primary and commercial cell lines and global ancestry was determined by standard estimates using single nucleotide variant-derived ancestry informative markers measured by whole genome sequence analyses.

| Cell line | Histology | Ancestry<br>(Standard<br>Estimate) |
| --- | --- | --- |
| ACI181 | Endometrioid | African |
| ACI80 | Endometrioid | African |
| NCIEC1 | Endometrioid | African |
| HEC1A | Adenocarcinoma NOS | East Asian |
| ISHIKAWA | Adenocarcinoma NOS | East Asian |
| SNGM | Adenocarcinoma NOS | East Asian |
| ACI52 | Endometrioid | European |
| ACI61 | Endometrioid | European |
| ACI68 | Endometrioid | European |
| AN3CA | Adenocarcinoma NOS | European |
| KLE | Adenocarcinoma NOS | European |
| MFE296 | Adenocarcinoma NOS | European |
| RL952 | Adenocarcinoma NOS | European |

### Quantitation of Peptide Ancestry Informative Markers (pAIMS) in Uterine Leiomyoma Tissues

We assessed pAIMs within multiplexed (TMT) quantitative proteomic analysis of formalin-fixed, paraffin embedded uterine leiomyoma tissues from self-described European-American (n=35) or African-American (n=31) women (Table 2, Discovery). We quantified 69,056 total peptides that included 33 high-confidence pAIMs (Supplemental Table 3). Of these candidates, 24 are of unknown clinical significance while 9 have been reported as not being pathogenic for disease (Supplemental Table 1A, ClinVar). Several of these candidates map to putative functional protein coding domains including intermediate filament domains within keratin 3 (R375G, >50% AF in Europeans) and keratin 13 (A187V, >90% in Europeans and East Asians) as well as within the dihydroxyacetone- (Dha-)binding domain of DHA kinase (A185T, >90% in Europeans and East Asians) (Supplemental Table 1A). We further assessed the performance of a subset of 18 pAIMs quantified within >50% of all uterine leiomyoma tissues to distinguish patient ancestry stratified by self-described race. Unsupervised hierarchical cluster analyses revealed that the abundance of pAIMs exhibiting high allele frequencies within European or African populations trended with self-described European or African-American race (Figure 5A). We also assessed performance of these candidates using a partial least squares (38) approach and identified, based on principle component analysis, these candidates served to explain 21% and 7% of the variance between African-American and European-American patient tissues (Figure 5B) and can further classify African-American from European-American patients with high accuracy (AUC=0.9834, P=1.6E^- 11^, Figure 5C). We also quantified pAIM abundance by multiplexed (TMT), quantitative proteomic analyses within an independent cohort of fresh-frozen uterine leiomyoma tissues from women of European (n=16) or African (n=10) ancestry confirmed using standard genotype estimates from companion WGS analyses, as recently described (39) (Table 2, Validation, Supplemental Table 4). We quantified a total of 61,283 total peptides among which 15 pAIMs were quantified in this cohort and 10 of which were co-quantified with discovery analyses (Supplemental Table 4). This validated subset was predominated by candidates of unknown clinical significance (ClinVar) and included a substitution in GC vitamin D-binding protein, i.e. (D451E, >50% Europeans). Assessment of pAIMs with allele frequencies >50% in African populations were significantly more abundant in African ancestry patients than pAIMS with allele frequencies of >50% in European ancestry populations (+1.0381 median fold difference, MWU p=0.003) (Figure 6A).

**Figure 5:**
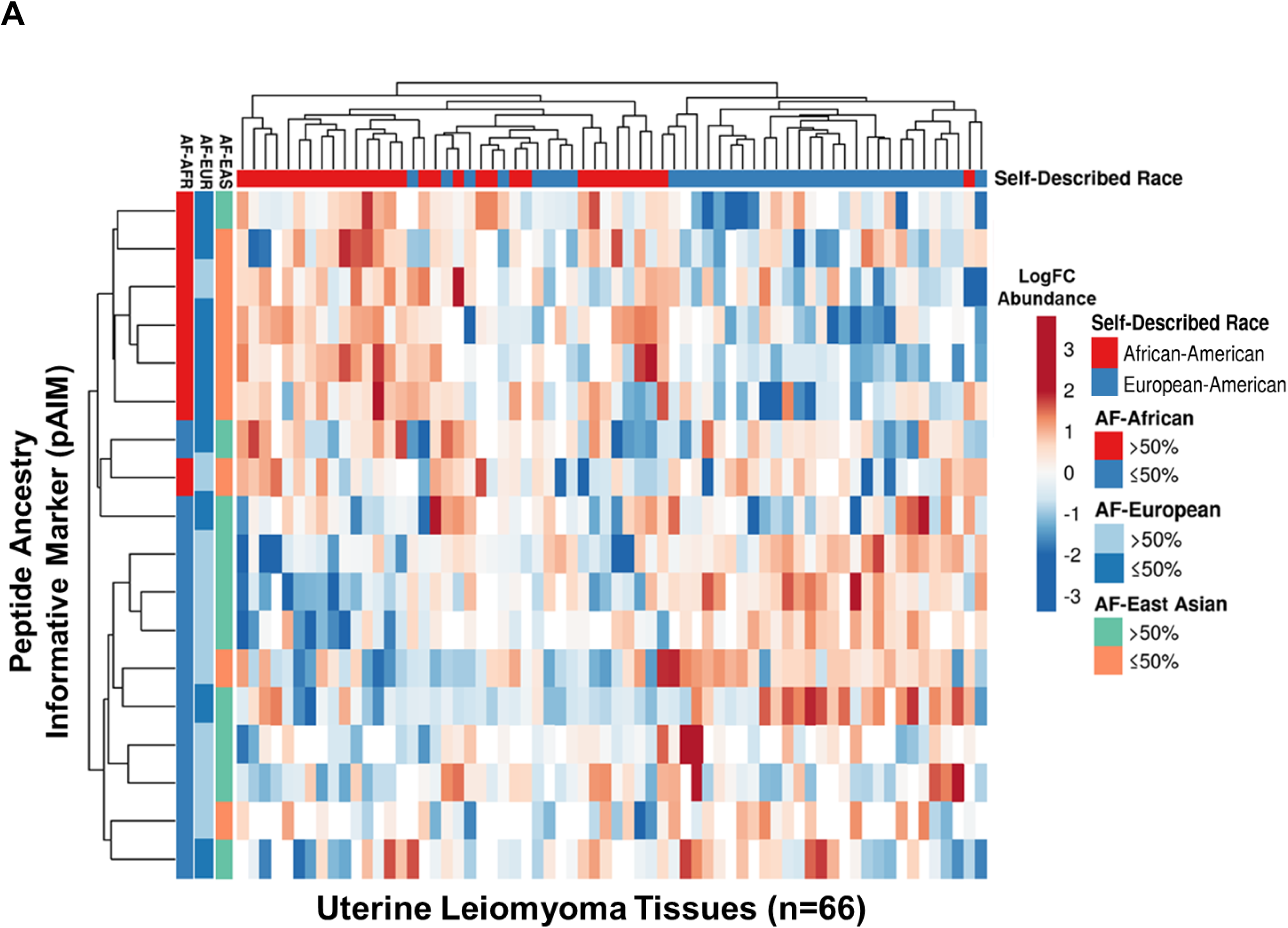
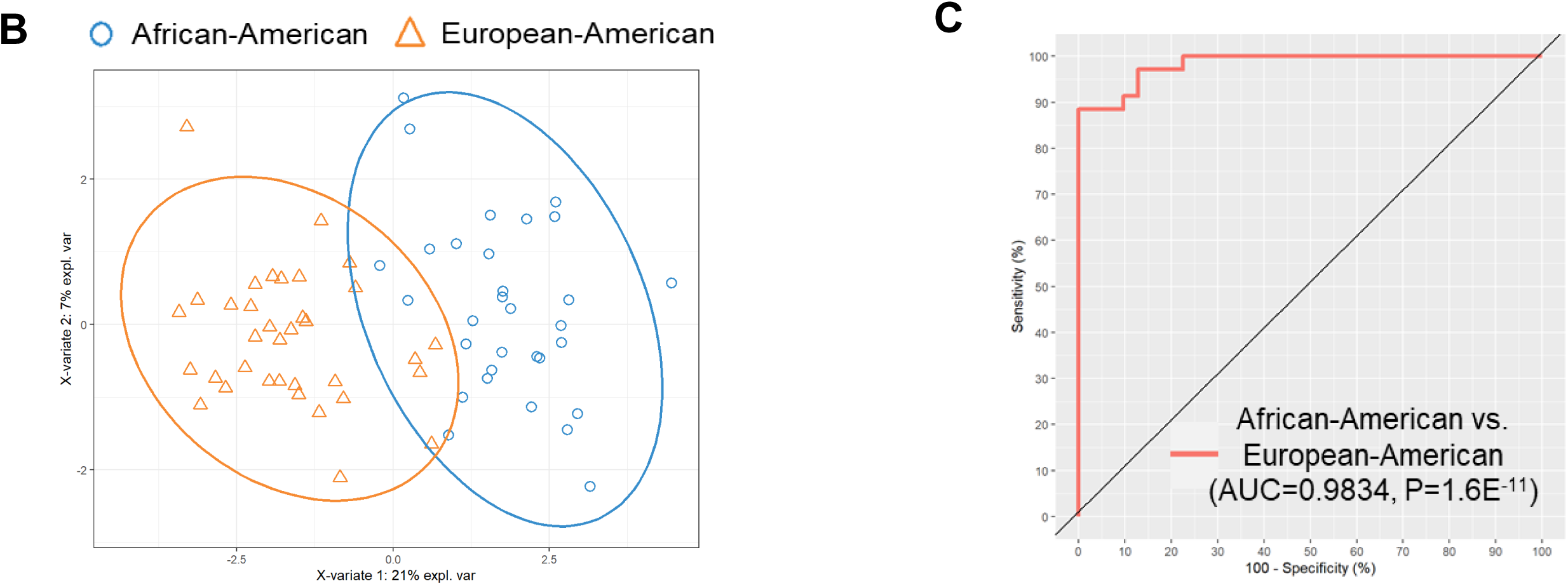
Quantitation of Peptide Ancestry Informative Markers (pAIMs) within Uterine Leiomyoma (ULM) Tissues Collected from a Cohort of Women of Self-Described Black or White Race. A: Data reflects unsupervised cluster analyses of 18 pAIMs quantified across n66 patient tissue samples; heatmap reflects clustering by Euclidean distance and complete linkage. B: Principle component analyses of 18 pAIMs, principal component 1 and principal component 2 that explain 21% and 7% of the total variance, respectively. C: Receiver operator curve assessing the performance of 18 pAIMs to classify African-American (n31) versus European-American (n35) patients.

**Figure 6:**
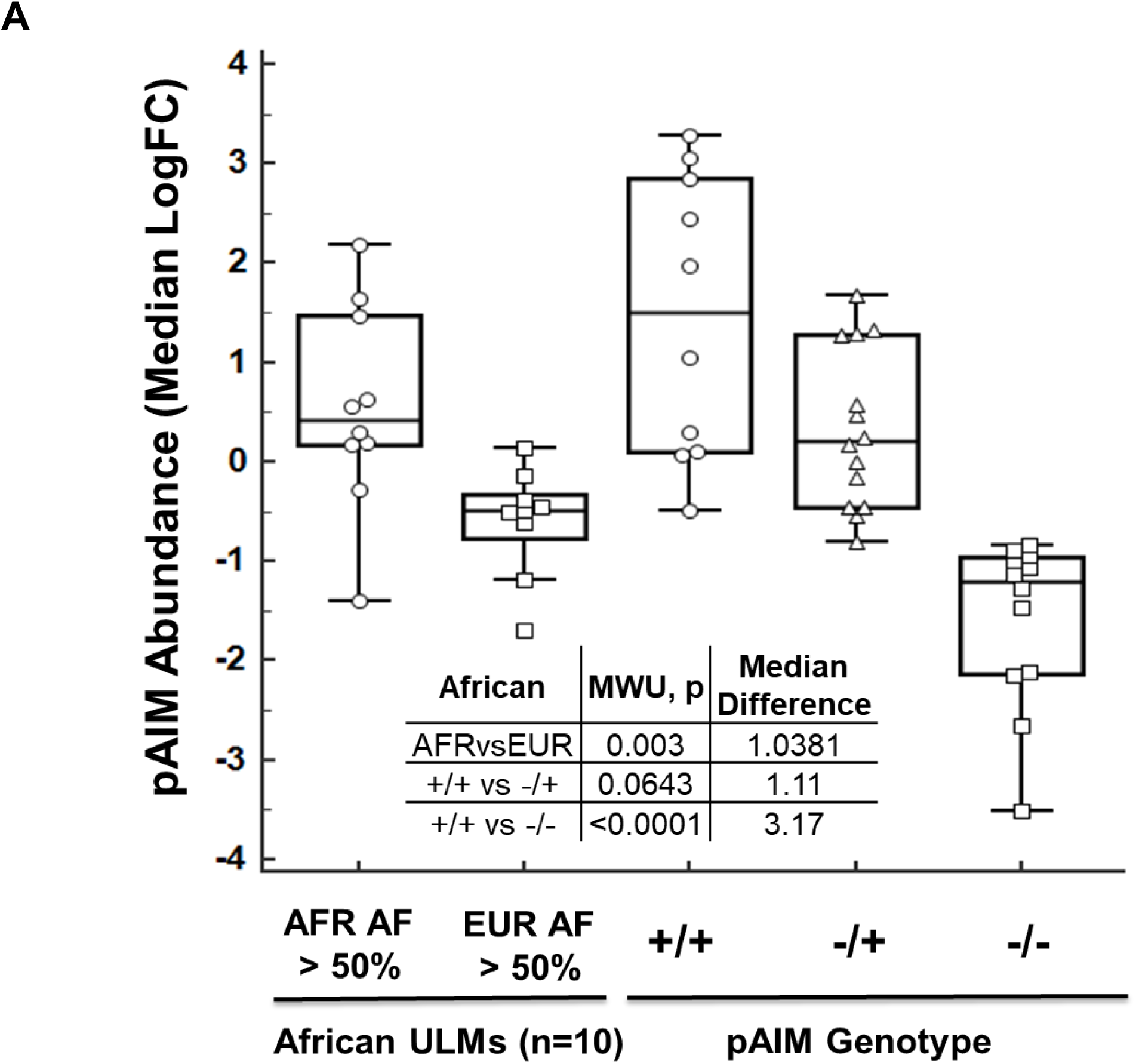
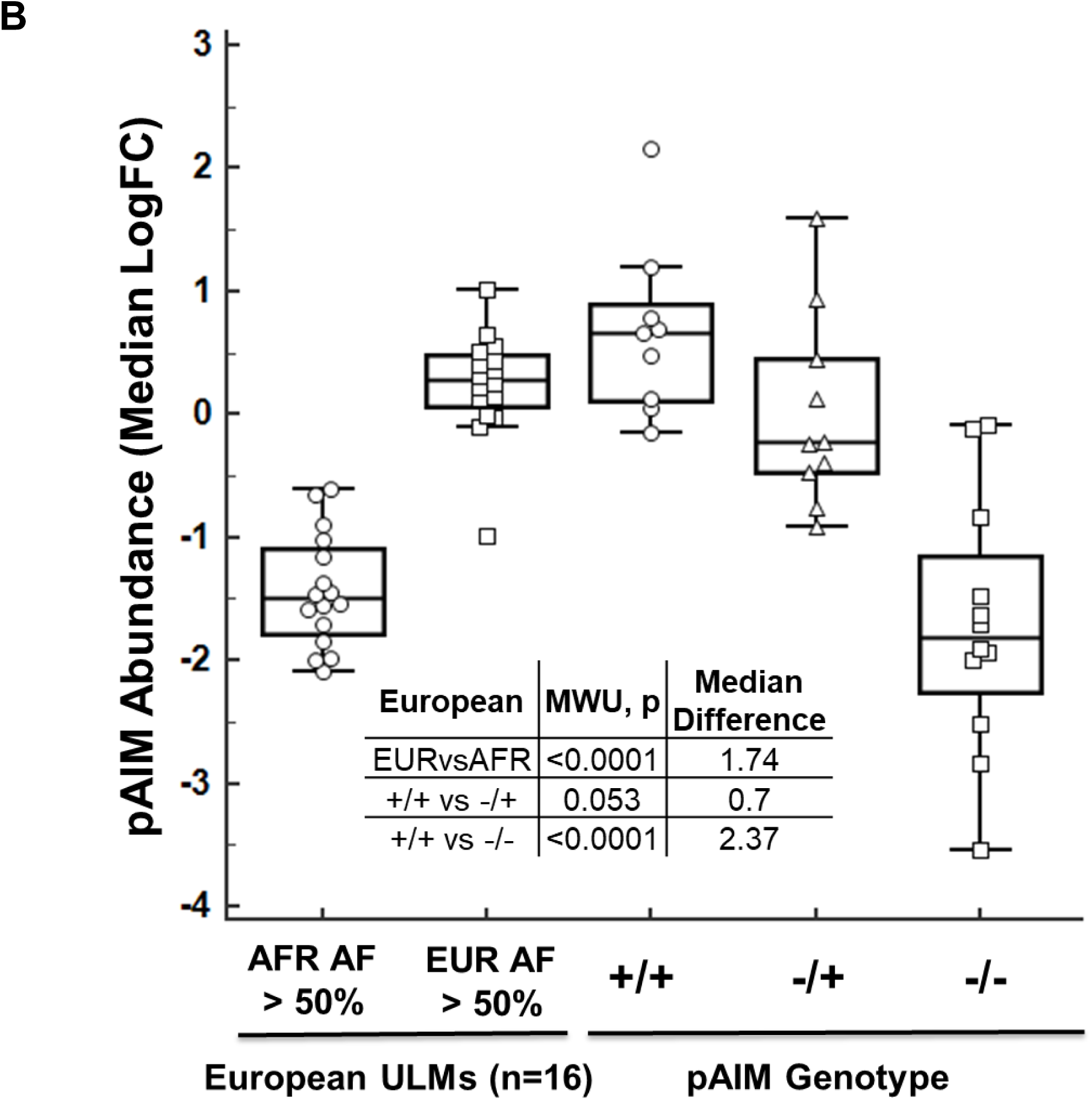
Validation of Peptide Ancestry Informative Markers (pAIMs) within Uterine Leiomyoma (ULM) Tissues Collected from Women with Global Ancestry determined by Standard Estimates. A: Box plots details the median logFC abundance for pAIMs exhibiting >50% allele frequencies in African (8 of 15 pAIMs quantified) reference populations across n10 ULMs sample from women with AFR ancestry as well as by pAIM variant allele genotype observed by companion whole genome sequence analyses. B: Box plots details the median logFC abundance pAIMs exhibiting >50% allele frequencies for European (5 of 15 quantified) reference populations quantified across n=16 ULMs sample from women with EUR ancestry as well as by pAIM variant allele genotype observed by companion whole genome sequence analyses.

**Table 2:** Uterine Leiomyoma (ULM) Tissues Analyzed by Quantitative Proteomic (Discovery and Validation) as well as Whole Genome Sequencing (Validation) Analysis. Global ancestry was determined by standard estimates for the validation cohort using single nucleotide variant-derived ancestry informative markers measured by whole genome sequence analyses.

| **Discovery - ULM**<br>**Tissues (n=66)** |  |
| --- | --- |
| **Ancestry**<br>**(Self-Described)** | **#** |
| European-American | 35 |
| African-American | 31 |

| **Validation - ULM**<br>**Tissues (n=26)** |  |
| --- | --- |
| **Ancestry**<br>**(Standard Estimate)** | **#** |
| European | 16 |
| African | 10 |

Conversely, we find that pAIMs with allele frequencies >50% in European ancestry populations were significantly more abundant in European ancestry patients than pAIMS with allele frequencies >50% in African populations (+1.74 median fold difference, MWU p<0.0001 (Figure 6B). Further, as observed in the endometrial cancer cell lines, we find that pAIMs abundances directly correlate with patients homozygous and heterozygous for the pAIM variant allele (+/+ and -/+ genotype) relative to those homozygous for the reference allele (-/-), i.e. African and European ancestry +/+ versus -/- patients (p<0.0001).

We further prioritized two pAIMs, i.e. a V237A substitution in protein Serpin Family A Member 1 (SERPINA1, K.DTEEEDFHVDQ(V/A)TTVK) with an allele frequency (AF) = 0.64 in African versus AF = 0.18 in European populations as well as an A19D substitution in Serine/threonine-protein phosphatase CPPED1 (CPPED1, R.TL(A/D)AFPAEK) with an AF = 0.75 in African versus AF = 0.15 in European populations for targeted analysis using a parallel reaction monitoring assay (40). Stable isotope standard (SIS) peptides were synthesized containing heavy isotope-labeled lysine residues and spiked into fibroid FFPE tissue digests collected from a cohort of African-American (n=3) and European-American women (n=3) previously assessed by multiplexed quantitative proteomic analysis (Supplemental Table 3). Quantitation of heavy SIS peptides revealed consistent abundance trends and high dot product similarity scores (dotp) (41) for the top five most abundant y-ions quantified for each peptide relative to library spectra across tissue digests from AA and EA women, i.e. 11.1% CV in abundance and average dotp = 0.92 ± 0.01 for the SERPINA1-V237A variant (Heavy SIS, Figure 7A, Supplemental Table 5) and 10.6% CV in abundance and average dotp = 0.91 ± 0.01 for the CPPED1-A19D variant (Heavy SIS, Figure 7B, Supplemental Table 5). Quantitation of endogenous SERPINA1-V237A and CPPED1-A19D peptides in tissue digests collected from African-American women from revealed high relative abundances and dot product similarity scores, i.e. SERPINA1-V237A peptide average dotp = 0.94 ± 0.01 (Figure 7A) and CPPED1-A19D peptide average dotp = 0.91 ± 0.03 (Figure 7B). Comparison of heavy SIS and endogenous peptide abundances further showed that SERPINA1-V237A pAIM variant was 6.1 ± 2.46 femtomole (fmol) and CPPED1-A19D variant 0.45 ± 0.12 fmol in fibroid tissue digests collected from African-American women (Table 3). Quantitative assessment of endogenous peptides in tissue digests collected from European-American women revealed peptide variants were largely below signal to noise or not detected and exhibited low dotp scores, i.e. SERPINA1-V237A average dotp = 0.33 ± 0.05 (Figure 7A) and CPPED1-A19D average dotp = 0.1 ± 0.15 (Figure 7B), suggesting these peptides were not present in these patient samples. Endogenous peptide abundance trends observed by targeted analysis are consistent with abundance trends quantified by multiplexed proteomic analysis for these patient tissue samples (Supplemental Table 3).

**Figure 7:**
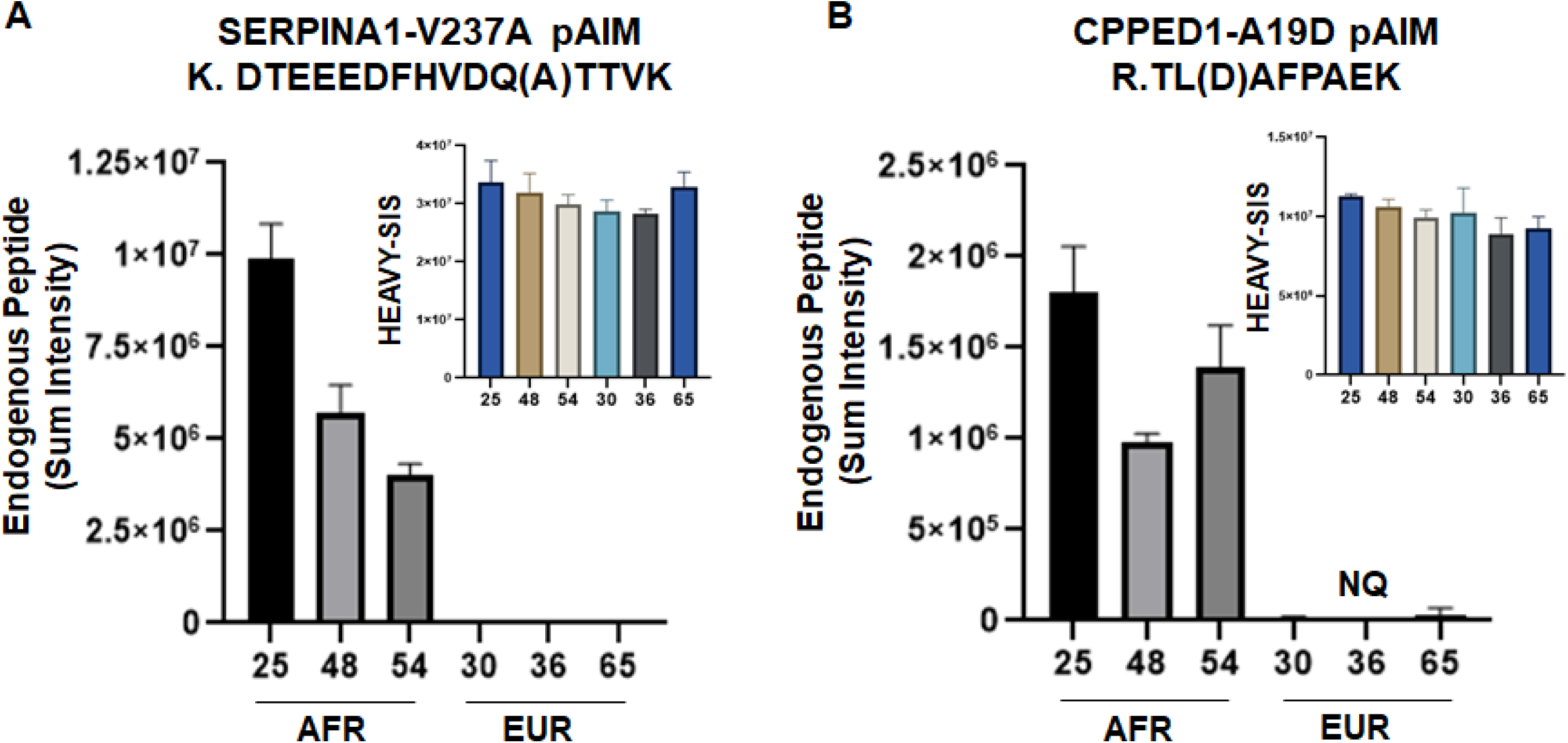
Targeted Quantitation of SERPINA1-V237A and CPPED1-A19D pAIM Candidates in Fibroid Tissue Digests from African-American and European-American Women. Stable isotope standard (SIS) peptides were synthesized with heavy-isotope labelled lysine residues corresponding to SERPINA1-V237A pAIM (K.DTEEEDFHVDQ(A)TTVK) and CPPED1-A19D (R.TL(D)AFPAEK) pAIM candidates and heavy SIS and endogenous peptide abundances were assessed by parallel reaction monitoring assay in fibroid FFPE tissue digests collected from African-American (AFR, n=3) and European-American (EUR, n=3) women. Data reflects sum intensity of the top five most abundant y-ions measured by peptide tandem mass spectrometry and triplicate, technical replicate injections. “NQ” = not quantified. A: Relative abundance of SERPINA1-V237A pAIM heavy SIS and endogenous peptides in AFR and EUR patient tissues. B: A: Relative abundance of CPPED1-A19D pAIM (R.TL(D)AFPAEK) candidates heavy SIS and endogenous peptides in AFR and EUR patient tissues.

**Table 3:** Targeted quantitation of SERPINA1-V237A and CPPED1-A19D stable isotope standard and endogenous pAIM peptide candidates in uterine fibroid tissue digests from African-American (AA) patients. fmol = femtomole.

| **Patient ID**<br>**& Self-**<br>**Described**<br>**Race** | **SERPINA1 - V237A**<br>**Endogenous/Heavy**<br>**SIS Peptide Ratio** | **SERPINA1 - V237A**<br>**Endogenous Peptide**<br>**Abundance (fmol)** | **CPPED1 – A19D**<br>**Endogenous/Heavy**<br>**SIS Peptide Ratio** | **CPPED1 – A19D**<br>**Endogenous Peptide**<br>**Abundance (fmol)** |
| --- | --- | --- | --- | --- |
| Patient<br>25_AA | 0.88 | 8.8 | 0.054 | 0.54 |
| Patient<br>48_AA | 0.54 | 5.4 | 0.03 | 0.3 |
| Patient<br>54_AA | 0.4 | 4.0 | 0.05 | 0.5 |

## Discussion

Comprehensive analysis of proteogenomic alterations correlating with patient ancestry are necessary to better understand the relationship between racial disparities underlying disease pathogenesis. In gynecologic malignancies, uterine neoplasms exhibit significant racial disparities, specifically increased incidence of uterine leiomyomas (42) as well as more aggressive endometrial cancers in African-American versus European-American patients (8,43,44). We and others have described race-specific molecular alterations correlating with disease outcome in endometrial cancers (45–47). As a foundational step to support our ongoing investigations of molecular alterations underlying racial disparities in gynecologic malignancies, we have investigated 1,037 nsSNPs exhibiting high allele frequencies within individuals of European, African, and East Asian populations that encode missense substitutions occurring within putative tryptic peptides, i.e. termed peptide ancestry informative markers (pAIMS) in cell line models and tissues derived from uterine neoplasms collected from women representing diverse ancestral backgrounds. Candidates prioritized were investigated *in silico* for their associations with disease as well as prevalence within putative functional protein domains and for predicted impact on protein function. We found that many pAIMs encode substitutions of unknown clinical significance (ClinVar), occur within functional protein domains and include subsets that are predicted to be deleterious to protein function. We further find that several pAIM variants are linked to an increased risk to develop cancer, such as the F31I substitution in aurora kinase A associated with colon cancer development in East Asian populations (16) and candidates that further track with cancer cell signaling, such as a substitution in nesprin-2 (SYNE2, H3309, rs8010699) that correlates with expression of cyclin-dependent kinase inhibitor 1 (CDK1/ p21) in HBV-related hepatocellular carcinomas (37). We also find that several candidates of unknown clinical significance occur within known functional protein domains including multiple immunoglobulin-like domains, such as in the major histocompatibility complex, class II, DP alpha 1 (HLA-DPA1) protein. These candidates further represent ancestrally-linked, variant peptides theoretically observable with routine LC-MS proteomic workflows encoding substitutions that may alter functional protein domain interactions as well as protein complex architecture.

Our efforts have quantified and validated pAIMs in uterine neoplasms from >100 women representing diverse ancestries and consist of a foundational set of ancestry-linked variant peptides observable by multiplexed proteomic analyses of uterine cells and tissues. Many pAIMs quantified are of unknown clinical significance in ClinVar and may warrant further investigation to better understand possible functional roles in the pathogenesis of uterine disease. Validated candidates were largely undocumented for disease pathogenesis in ClinVar but were associated with altered biological functions. One such candidate encodes a substitution in GC vitamin D-binding protein, i.e. (D451E, rs7041), a protein that is important in regulating transport of vitamin D, exhibiting a higher allele frequency within individuals of European (58%) versus African (7.3%) descent. rs7041 has been shown to correlate with altered levels of GC within European and African-Americans (48) and further to impact vitamin D metabolism in pregnant women (49). Comparison of cell line and tissue analyses revealed quantitation of pAIMs encoded within Protein-L-isoaspartate(D-aspartate) O-methyltransferase (V178I, >80% AF in African and East Asians), Serine/threonine-protein phosphatase CPPED1 (A19D, >75% AF in Africans) and AH receptor-interacting protein (Q228K, >99% AF in European and East Asians) across all samples. Protein-L-isoaspartate(D-aspartate) O-methyltransferase (PCMT1) regulates methyl esterification of L-isoaspartyl and D-aspartyl residues and has been shown to repair damage to aspartate resides that can occur in an age-dependent manner (50). Interestingly, the I178 variant of PCMT1 has been shown to result in increased catalytic activity and a more thermostable version of PCMT1, while the V178 variants exhibits greater substrate affinity. Serine/threonine-protein phosphatase CPPED1 has been shown to function as a tumor suppressor in bladder cancer via inactivation of Protein Kinase B (AKT) through dephosphorylation of S473 leading to inhibition of cell cycle progression and promotion of apoptosis (51). Although the A19D ancestry-linked substitution we have observed is not predicted to impact CPPED1 function, we have previously reported higher mutational frequencies in the gene encoding tumor suppressor phosphatase and tensin homolog (PTEN) in European-American versus African-American endometrial cancer patients, a regulator of AKT signaling, that correlates with improved outcome in European-American women (46). This intersection suggests that regulation of AKT signaling associated protein machinery may be linked to patient ancestry and warrants further investigation of these relationships and their impact on disease pathogenesis.

Our analyses provide proof of concept that estimates of European, African and East Asian ancestry can be determined by routine proteomic analysis of disease-relevant patient tissues. We find that pAIMs exhibiting higher allele frequencies within European, African or East Asian ancestry individuals are quantified at significantly greater abundances in cell lines and tissue samples derived from individuals of similar self-described ancestry, i.e. European-American or African-American, and further exhibited abundance trends correlating with patient genotype. Our findings also underscore that quantitation of pAIMs is feasible in formalin-fixed, paraffin embedded as well as fresh frozen human tissue samples. These data extend previous efforts showing that genetically variant peptides can provide estimates of global ancestry in human hair and bone and underscore that these estimates can be determined using multiplexed proteomic data similar to that being generated for population-scale proteogenomic studies focused on improving our understanding of cancer. We further show that we can accurately estimate global ancestry by quantifying as few as 20 population-specific pAIMs in comparison with genotype analyses of ∼266K genetic markers using standard estimates (R^2^>0.99). Although our focus has been to characterize pAIMs in uterine neoplasms, these candidates are not tissue dependent and represent a generalized set of markers for proteoancestry assessment in diverse tissues and cell lines. To that end, comparison of proteins encoding pAIMs with large-scale proteomic analyses of diverse human cell and tissue types (52) suggests that many of these protein targets are ubiquitously expressed across various tissues, including diverse immune cell types (data not shown) and that quantitation of pAIMs in a wide range of biological samples is likely possible. One limitation of this study we further note is that several pAIMs quantified also map to largely predicted proteins (Uniprot.org and noted in Supplemental Tables 2 thru 4) underscoring the need to further refine pAIMs panels with the goal of prioritizing candidates that estimate patient ancestry with the greatest sensitivity and specificity for a given organ site. To this end, we further provide proof of concept data for the targeted quantitation of two pAIM candidates, i.e. SERPINA1-V237A and CPPED1-A19D that exhibit higher allele frequencies with African versus European populations in fibroid tissue digests from African-American and European-American women. These analyses verified the relative abundance trends observed for these candidates by multiplexed, quantitative proteomic analysis and further showed these candidates exhibited elevated abundance in African-American women, but were largely not detected within European-American women, consistent with the known allele frequencies for these variants in these populations. These findings will support future efforts focused on optimizing tissue-specific, stable isotope-labelled pAIMs panels that can be co-quantified during primary clinical sample analyses.

We have characterized ancestry-linked peptide variants in endometrial cancer cell lines and patient tissues representing a foundational step towards investigating relationships linking patient ancestry with the proteome, genome and disease pathogenesis. To this end, recent investigations have revealed that clinical serum biomarkers can exhibit altered performance directly as a product of differences in patient genetic ancestry, underscoring the need to integrate proteomics with genetic ancestry assessments at the level of investigational analyses such as biomarker discovery (53). Racial disparities in the incidence and outcome of uterine neoplasms persists following adjustments for socioeconomic variables, such as patient access to care (54) and has fueled efforts to define ancestry-linked molecular alterations that may drive these differences. Our ability to discern these relationships will emerge through the assessment of more ancestrally-diverse populations of women suffering from gynecologic disease as well as within disease-relevant *in vitro* and *in vivo* models. We recently showed that historic cell line models of cancer have been predominantly established from individuals of European ancestry (55), noting that cancers impacted by racial disparities, such as endometrial cancer, often lacked cell line models from non-European individuals. We confirm here three endometrial cancer cell line models, i.e. NCI-EC1, ACI-80 and ACI-181 are derived from women of African ancestry representing *in vitro* models to investigate ancestry-linked disease biology in endometrial cancers. This work will support ongoing efforts focused on linking the proteome and genome with patient ancestry to better understand relationships with disease mechanisms driving racial disparities in the pathogenesis of uterine neoplasms.

## Methods

### Prioritization and *In Silico* Analyses of Peptide Ancestry Information Markers

80,855,907 SNPs for European (i.e. FIN, GBR, IBS, and TSI; N=404), African (i.e. ESN, GWD, LWK, MSL, and YRI; N=504), and East Asian (i.e. CDX, CHB, CHS, and KHV; N=400) were downloaded from the 1000 Genomes Project (23). SNPs were further mapped to refSeq version 85 to identify candidate missense variants (646,768 nsSNPs) occurring within putative tryptic peptides ≥6 or <40 amino acids in length that did not encode isoleucine to leucine substitutions and were further filtered for those exhibiting ≥ 50% difference in allele frequency between European, African, and East Asian populations. These substitutions were then mapped to several RefSeq proteome databases to confirm amino acid substitution positions and tryptic specificity resulting in a final subset of 1,037 candidates. nsSNPs encoding pAIMs were mapped to rsIDs using Kaviar (http://db.systemsbiology.net/kaviar/cgi-pub/Kaviar.pl) against hg19 (GRCh37). Candidates were further assessed using the Variant Effect Predictor tool (https://useast.ensembl.org/Homo_sapiens/Tools/VEP) against hg19 (GRCh37) and extracted metadata detailing putative clinical significance, i.e. ClinVar traits and clinical significance, as well as predicted impact on protein function, i.e. SIFT (28) and Poly-Phen-2 (29) and PROVEAN (30) outputs. Candidates were further mapped to the Uniprot resource and amino acid position of pAIM substitutions were correlated with putative functional domains within parent proteins and pAIM substitutions positions mapping with putative domain regions were prioritized for downstream analyses. Functional enrichment analyses was performed using the “express analysis” settings in MetaScape (56). Determination of fixation index for pAIMs. Fixation index (FST) values were calculated using reference and alternative allele frequency data from the 1000 genomes project. The following equation was used; FST = (HT-HS)/HT, where HT is the expected heterozygosity in the total population and HS is the expected heterozygosity in the subpopulation (57).

### Random Sampling Analysis

After the 1037 pAIMs were selected from the 1000 Genomes Project training data, we then performed ADMIXTURE (34) randomly sampling 20 pAIMs without replacement, using the supervised learning version (33) with the training data as fixed for their respective ancestries (K=3 for African, European, and East Asian). Test data also came from the following 1000 Genomes Project populations (23), which were not included in the training: JPT to represent East Asian, CEU to represent European, and both ACB and ASW as admixed African ancestry. This procedure was repeated 500 times with the mean and standard deviation calculated and plotted against the estimates for these same samples from 266,403 SNPs. This larger set of sites was selected in the standard way (58), where they were LD pruned (using the PLINK(59) command --indep-pairwise 50 5 0.1), and filtered for allele frequency of 5%.

### Sample Prep for Proteomics

#### Cell Lines

Commercial endometrial cancer cell lines, i.e. MFE-296 cells were purchased from DSMZ GmbH (Berlin, Germany) and cultured in MEM with 10% fetal bovine serum (FBS) and 1X penicillin/streptomycin (Pen/Strep). SNG-M cells were purchased from the JCRB cell bank (Osaka, Japan) and cultured in Ham’s F12 median with 10% FBS and 1X Pen/Strep. HEC1A (McCoy’s 5a with 10% FBS and 1X Pen/Strep), RL95-2 (DMEM:F12, 0.005 mg/mL insulin, 10% FBS and 1X Pen-Strep), Ishikawa (MEM with 2mM Glutamine, 5% Fetal Bovine Serum (FBS) and 1X Pen/Strep), AN3CA (EMEM with 10% FBS and 1X Pen/Strep) and KLE (DMEM:F12 with 10% FBS and 1X Pen/Strep) cells were from ATCC (Manassas, VA). NCI-EC1, ACI-80, ACI-181, ACI-52, ACI-61 and ACI-68 were obtained from John Risinger at Michigan State University (East Lansing, MI) and maintained in DMEM:F12 with 10% FBS and 1X Pen/Strep (ATCC). Cell lines were plated at equivalent densities and lysed at ∼80% confluency in 1% SDS and 10mM Tris-HCl, pH 7.4. Cell lysates (30 µg) were run approximately one inch into a 4-15% bis-acrylamide gel (Bio-Rad) and processed for in-gel digestion as previously described (60). Peptide extracts were labelled with TMT-11 reagents (Thermo Fisher Scientific) as described below.

#### Uterine Leiomyoma Tissues (Discovery cohort)

Age-matched, formalin-fixed, paraffin embedded uterine leiomyomas from self-described white and black women were obtained under an institutional-review board approved protocol from Inova Fairfax Hospital (Falls Church, VA). Tissue specimens were hematoxylin and eosin (H&E) stained and evaluated by a pathologist to confirm diagnosis. Thin (10 µm) tissue sections were cut using a cryostat and placed on polyethylene naphthalate membrane slides. After staining with aqueous H&E, laser microdissection (LMD) was used to harvest tumor cells from thin sections.

#### Uterine Fibroid Tissues (Validation cohort)

Flash-frozen uterine leiomyomas were obtained under an institutional-review board approved protocol from Johns Hopkins Memorial Institute (Baltimore, MD). Tissues were cryopulverized, resuspended in 100 mM triethylammonium bicarbonate (TEAB), and sonicated. Protein was quantified by BCA Protein Assay (Thermo Scientific) and 50 µg of total protein in 100 mM TEAB and 10% acetonitrile was incubated at 99°C for 1 h.

### Sample digestion, TMT labeling and offline fractionation

Sample digestion, TMT labeling and offline fractionation of tissue samples was performed as recently described (61). Briefly, tissues were collected in 20 µL of 100 mM TEAB in MicroTubes (Pressure Biosciences, Inc). Following the addition of 1 µg of SMART Digest Trypsin (Thermo Scientific) to each sample, MicroTubes were capped with MicroPestles. Pressure-assisted lysis and digestion was performed in a barocycler (2320 EXT, Pressure BioSciences, Inc) by sequentially cycling between 45 kpsi and atmospheric pressure for 60 cycles at 50 °C. The peptide digests were transferred to 0.5 mL microcentrifuge tubes, vacuum-dried, resuspended in 100 mM TEAB, pH 8.0 and the peptide concentration of each digest was determined using the bicinchoninic acid assay (BCA assay). Forty-fifty micrograms of peptide from each sample, along with a pooled reference sample assembled from equivalent amounts of peptide digests pooled from individual patient samples for individual sample sets, were aliquoted into a final volume of 100 µL of 100 mM TEAB and labeled with tandem-mass tag (TMT) isobaric labels (TMT10 or 11plex™ Isobaric Label Reagent Set, Thermo Fisher Scientific) according to the manufacturer’s protocol. Each TMT sample plex was fractionated by basic reversed-phase liquid chromatography (bRPLC) into 96 fractions through development of a linear gradient of acetonitrile (0.69%/min). Concatenated fractions (36 total pooled samples for the endometrial cancer cell lines, FFPE ULMs, 24 total for the frozen ULMs) were generated in a serpentine fashion for global LC-MS/MS analysis.

### Quantitative Proteomics - LC-MS/MS Analyses

The TMT sample multiplex bRPLC fractions were resuspended in 100 mM NH_4_HCO_3_ and analyzed by LC-MS/MS employing a nanoflow LC system (EASY-nLC 1200, Thermo Fisher Scientific) coupled online with an Orbitrap Fusion Lumos MS or Q-Exactive HF-X (Thermo Fisher Scientific). In brief, each fraction (∼500 ng total peptide) was loaded on a nanoflow HPLC system fitted with a reversed-phase trap column (Acclaim PepMap100 C18, 20 mm, nanoViper, Thermo Scientific) and a heated (50 °C) reversed-phase analytical column (Acclaim PepMap RSLC C18, 2 µm, 100 Å, 75 µm × 500 mm, nanoViper, Thermo Fisher Scientific) coupled online with the MS. Peptides were eluted by developing a linear gradient of 2% mobile phase B (95% acetonitrile, 0.1% formic acid) to 32% mobile phase B over 120 min at a constant flow rate of 250 nL/min. For both instrument platforms, the electrospray source capillary voltage and temperature were set at 2.0 kV and 275 °C, respectively. High resolution (R=60,000 at m/z 200) broadband (m/z 400-1600) mass spectra (MS) were acquired, followed by selection of the top 12 most intense molecular ions in each MS scan for high-energy collisional dissociation (HCD). Instrument specific parameters were set as follows for each instrument platform. Orbitrap Fusion Lumos - Full MS: AGC, 5e5; RF Lens, 30%; Max IT, 50ms; Charge State, 2-4; Dynamic Exclusion, 10ppm/20 sec; MS2: AGC, 1e5; Max IT, 120ms; Resolution, 50k; Quadrupole Isolation, 0.8 m/z; Isolation Offset, 0.2 m/z; HCD, 38; First Mass, 100. Q Exactive HF-X - Full MS: AGC, 3e6; RF Lens, 40%; Max IT, 45ms; Charge State, 2-4; Dynamic Exclusion, 10ppm/20 sec; MS2: AGC, 1e5; Max IT, 95ms; Resolution, 45k; Quadrupole Isolation, 1.0 m/z; Isolation Offset, 0.2 m/z; NCE, 34; First Mass, 100; Intensity Threshold, 2e5; TMT Optimization, On.

### DNA extraction for endometrial cancer cell lines and uterine fibroid tissue samples

Cells were lysed in SNET Digestion buffer (20 mM Tris, 5 mM EDTA, 400 mM NaCl, 1% w/v SDS) with Proteinase K (Thermo Fisher Scientific, Pittsburgh, PA) overnight at 55 °C with intermittent shaking. An equal volume of 25:24:1 phenol:chloroform:isoamyl alcohol (Thermo Fisher Scientific, Pittsburgh, PA) was added, mixed by inversion and centrifuged at 21,000 x *g* for 5 min. The supernatant was removed to a new tube, half sample volume of 7.5 M ammonium acetate and one and a half sample volumes of 100% ethanol were added, mixed by inversion and centrifuged at 14,000 x *g* for 10 min. The DNA pellet was washed with 70% ethanol and centrifuged at 14,000 x *g* for 5 min. The supernatant was removed and the pellet was air-dried and resuspended in 50 μL of 10 mM Tris buffer. Tissue scrolls were generated from OCT-embedded fresh-frozen tumors and collected in ATL buffer (Qiagen Sciences LLC, Germantown, MD) followed by storage at -80 °C until isolation. Samples were normalized to 360 μL ATL buffer, 40 μL of Proteinase K was added for lysis and incubated at 56 °C for 4 h with intermittent shaking. Isolation was performed according to the manufacturer’s protocol (DNA Purification from Tissues) using the QiAamp DNA Mini Kit (Qiagen Sciences LLC, Germantown, MD). DNA was eluted after a 10 min incubation with 40 μL of Buffer AE, followed by another 10 min incubation with 160 μL of nuclease-free water (Thermo Fisher Scientific) and reduced to 50 μL using a CentriVap Concentrator (Labconco, Kansas City, MO). Quantity and 260/280 purity reading was established using the Nanodrop 2000 Spectrophotometer (Thermo Fisher Scientific) and Quant-iT PicoGreen dsDNA Assay Kit (Thermo Fisher Scientific) according to manufacturer’s protocols using eight standards from 0 to 50 ng/uL. Samples were run in triplicate and measurements were taken on a SpectraMax M4 microplate reader (Molecular Devices, San Jose, CA).

### DNA PCR-free library preparation and whole genome sequencing

TruSeq DNA PCR-free Library Preparation Kit (Illumina, San Diego, CA) was performed following manufacturer’s instructions. Briefly, genomic DNA (gDNA) was diluted to 20 ng/μL using Resuspension Buffer (RSB, Illumina) and 55 μL were transferred to Covaris microTubes (Covaris, Woburn, MA). The normalized gDNA was then sheared on an LE220 focused-ultrasonication system (Covaris) to achieve target peak of 450 bp with an Average Power of 81.0 W (SonoLab settings: duty factor, 18.0%; peak incident power, 45.0 watts; 200 cycles per burst; treatment duration, 60 s; water bath temperature, 5 °C – 8.5 °C). The quality of the final DNA libraries was assessed with the High Sensitivity dsDNA (AATI). Per manufacturer’s protocol, library peak size was in the range of 550 to 620 bp. The DNA libraries were quantified by real-time quantitative PCR, using the KAPA SYBR FAST Library Quantification Kit (KAPA Biosystems, Boston, MA) optimized for the Roche LightCycler 480 instrument (Roche). DNA libraries were then normalized to 2 nM and clustered on the Illumina cBot 2 at 200pM using a HiSeq X Flowcell v2 and the HiSeq X HD Paired-End Cluster Generation Kit v2. Paired-end sequencing was performed with the HiSeq X HD SBS Kit (300 cycles) on the Illumina HiSeq X.

### Data Processing – Proteomics

Peptide ancestry informative marker identifications are generated by searching .RAW data files with a publicly-available human proteome database (RefSeq human proteome, downloaded 9/22/17 and updated with entries from 8/19/2019 when pAIM amino acid positions did not map to entries in 9/22/17 version) appended with 1,037 full-length proteins encoding predicted pAIM substitutions of interested using Mascot (v2.6.0, Matrix Science) and Proteome Discoverer (v2.2.0.388, Thermo Fisher Scientific) as previously described (61). Samples are searched using the following parameters: precursor mass tolerance of 10 ppm, fragment ion tolerance of 0.05 Da, a maximum of two tryptic miscleavages, dynamic modifications for oxidation (15.9949 Da) on methionine residues and TMT reporter ion tags (229.1629 Da) on peptide N-termini and lysine residues. Peptide spectral matches (PSMs) are filtered using a false-discovery rate (FDR) < 1.0% (Percolator q-value < 0.01). TMT reporter ion intensities are extracted at a mass tolerance of 20 ppm and PSMs lacking TMT reporter ion signal in the pooled reference channel, in all patient TMT channels or exhibiting an isolation interference of ≥50% were excluded from downstream analyses. Normalized peptide abundance was determined by calculating TMT reporter ion ratios relative to a pooled reference channel and applying a mode centered, z-score transformation of each sample channel per TMT multiplex, i.e. normalized (PSM (Log_2_Ratio) = (PSM (Log_2_Ratio) – ModeCenter (PSM (Log_2_Ratio) / σ (PSM (Log_2_Ratio). Peptide fragment ion spectra for pAIM peptide variants are confirmed to encode the substitution of interest by automated inspection of MS2 fragment ion spectra in .mzML converted .RAW files (MSConvert, Proteowizard) using SpectrumAI (36) and the following settings, fragment ion tolerance was set to 50 ppm, relative set to True and a candidate was considered high-confidence if diagnostic ions flanking the substitution of interest were identified in at least one sample multiplex in endometrial cancer cell line and uterine leiomyoma cohorts. Differential analyses of pAIM abundance was performed using Mann-Whitney U rank sum testing in MedCalc (version 19.0.3). pAIMs variants were visualized in heatmaps and by principle component analysis (PCA) using default settings in the ClustVis web tool (https://biit.cs.ut.ee/clustvis/). Sparse Partial Least Squares Discriminant analysis (sPLS-DA) (38) was performed using mixOmics (ver 6.8.5) in RStudio (ver 3.6.0). The sPLS-DA model was run in regression mode for an optimized 18 feature set for two principal components. Plot loadings were calculated using the median to assess contribution onto the first principal component. The AUC and ROC curve were generated from the sPLS-DA model.

### Parallel reaction monitoring analysis of SERPINA1-V237A and CPPED1-A19D pAIM candidates

Stable isotope standard versions of SERPINA1-V237A (DTEEEDFHVDQ(A)TTVK) and CPPED1-A19D (TL(D)AFPAEK) pAIM peptide candidates were synthesized with heavy isotope labelled lysine (K, i.e. C13(6), 15N(2) residues (Heavy SIS peptides, Thermo Fisher Scientific). A total of 10fmol final of heavy SIS peptides was spiked into ∼1 ug total fibroid tissue digest for each patient sample and samples were analyzed in triplicate by LC-MS/MS employing a nanoflow LC system (EASY-nLC 1200, Thermo Fisher Scientific) coupled online with a Q-Exactive HF-X (Thermo Fisher Scientific). Briefly, each sample was loaded on a nanoflow HPLC system fitted with a reversed-phase trap column (Acclaim PepMap100 C18, 20 mm, nanoViper, Thermo Scientific) and a heated (50 °C) reversed-phase analytical column (Acclaim PepMap RSLC C18, 2 µm, 100 Å, 75 µm × 150 mm, nanoViper, Thermo Scientific) coupled online with the MS. Peptides were eluted by developing a linear gradient of 2% mobile phase B (95% acetonitrile, 0.1% formic acid) to 32% mobile phase B over 60 min at a constant flow rate of 300 nL/min, then to 99% mobile phase B over an additional 15 min, followed by flushing and re-equilibration prior to the next injection. Source capillary voltage and temperature were set at 2.0 kV and 275 °C, respectively. High resolution (R=120,000 at m/z 200) broadband (m/z 400-1400) mass spectra (MS) were acquired in profile mode, followed by selection of the top 12 most intense molecular ions in each MS scan for high-energy collisional dissociation (HCD). Instrument specific global parameters were set as follows: Full MS: AGC, 3e6; RF Lens, 40%; Max IT, 50 ms; Charge State, 2-3; MS2: AGC, 1e5; Max IT, 50 ms; Resolution, 15k; Quadrupole Isolation, 1.0 m/z; Isolation Offset, 0.2 m/z; NCE, 30; Intensity Threshold, 1.6e5; Dynamic Exclusion, Off; TMT Optimization: Off. Inclusion list entries consisted of the endogenous/variant and heavy/variant SIS pAIMs peptides and the top 12 most abundance molecular ions were selected for data-dependent acquisition when not monitoring inclusion list masses. All targeted peptides were fragmented at an NCE of 28. RAW data files were searched using parameters noted above excluding static modifications for TMT reporter ion tags. RAW data files and Proteome Discoverer search results were imported into Skyline-daily (64-bit, v. 21.0.9.118) (62) using the “Import PRM Peptide Search” workflow and the top five y-ions quantified for heavy stable isotope standard (SIS) and endogenous versions of the SERPINA1-V237A (z = 3+) and CPPED1-A19D (z = 2+) pAIM peptide candidates were extracted using default settings for orbitrap instrumentation. Sum y-ion intensities and dot product similarity scores were exported from Skyline and plotted using Graphpad Prism (v 8.4.3).

### DNA WGS processing and variant calling

WGS sample raw reads were aligned to the hg19 reference genome and further processed through the Resequencing workflow within Illumina’s HiSeq Analysis Software (HAS; Isis version 2.5.55.1311; https://support.illumina.com/sequencing/sequencing_software/hiseq-analysis-software-v2-1.html). This workflow utilizes the Isaac read aligner (iSAAC-SAAC00776.15.01.27) and variant caller (starka-2.1.4.2) (63), the Manta structural variant caller (version 0.23.1) (64), and the Canvas CNV caller (version 1.1.0.5) (65).).

### Global ancestry proportion of EC cell lines

To estimate admixture proportions, we used reference samples with known ancestry from the 1000 Genome Project (23). These reference samples were comprised of 99 European samples from the CEU population, 108 African samples from the YRI population, and 208 Asian samples from the CHB and JPT populations (414 total references samples). Working with genotype data from these reference samples and each of the endometrial cancer cell lines, we first used bcftools(66) to remove Indels and non-biallelic variants. We then used Plink v1.90b3.32 (59)to remove singleton sites (i.e. variants with only one alt allele copy in the dataset. After this processing, we merged these datasets with Plink by taking the intersect of autosomal variants and removing any ambiguous A/T and G/C variants from this combined data. We then pruned for linkage using the plink linkage pruning algorithm command of --indep-pairwise 50 5 0.5, which uses a window of 50 with an r^2^ greater than 0.5 and a SNP step of 5. Ancestry was estimated for each cell line independently, and so this process was repeated independently for each cell line. At the end of this processing, we were left with ∼500,000 LD-pruned variants per reference cell line merged dataset (range 459,592 – 701,192).

### Global ancestry proportion of uterine fibroid patients

We predicted sample super population ancestries using the methods implemented in Peddy (67). Briefly, principal component reduction was performed on genotype calls at specific loci from 2,504 samples in the 1000 Genomes project and a support vector machine (SVM) classifier was trained on the resulting first four components, using known ancestries as the training labels. Genotype calls at the same loci from each sample collected in this study were then mapped to principal component space and the trained SVM was used to predict ancestries. All classifier prediction probabilities were >0.89.

### Data Availability

Supplemental data tables include chromosomal loci, missense substitutions and amino acid position, reference SNP Ids (rsIDs) corresponding to peptide ancestry informative markers (pAIMs), as well as pAIM abundances quantified in endometrial cancer cell lines and uterine leiomyoma tissues. RAW LC-MS data files will be deposited in ProteomeXchange.

## Acknowledgments

The authors would like to thank Tracy Litzi, BS, Glenn Gist, BS, Julie Oliver, MS, Dave Mitchell, MS, Domenic Tommarello, MS and Wei Ao, BS for their technical contributions as well as Albert Dobi, PhD, Gyorgi Petrovics, PhD and Sara Scannell, BS for critical review of this manuscript. GLM is a consultant for Kiyatec, GSK, and Merck. TPC is a ThermoFisher Scientific, Inc SAB member and receives research funding from AbbVie. GJP has received a patent based on concepts presented in this study, (US 8,877,455 B2, Australian Patent 2011229918, Canadian Patent CA 2794248, and European Patent EP11759843.3). Disclaimer: The views expressed herein are those of the authors and do not reflect the official policy of the Department of Army/Navy/Air Force, Department of Defense, or U.S. Government.

## Author Contributions

Contributed to conception: NWB, TDO, TPC, GLM. Contributed to experimental design: NWB, TDO, TPC. Contributed to data acquisition, analysis and/or interpretation of data: NWB, TDO, CMT, TA, BLH, KC, MZ, PT, ARS, AJ, ZG, JL, CT, CD, MW. Drafted and/or revised the article: NWB, TDO, ARS, MK, CDS, HH, MC, GJP, JS, AAH, JRR, KD, YC, TPC, GLM. Acquired funding for the research: YC, GLM. All authors read and approved the final manuscript.

## Supplemental Tables

**Supplemental Table 1A** – Peptide Ancestry Informative Markers. Column headers denote the following ProteinAccession_pAIM: Refseq protein accession and missense substitution and amino acid position encoded by the nonsynonymous single nucleotide variant encoding pAIM, SNP-ID: chromosome and genomic loci (hg19), GeneSymbol: Gene name, rsID: reference SNP, VAR_Position: amino acid and position of the reference and variant, AF-African: pAIM allele frequency in African reference populations (1000 genomes project), AF-European: pAIM allele frequency in European reference populations (1000 genomes project), AF-EastAsian: pAIM allele frequency in East Asian reference populations (1000 genomes project), SIFT-D: YES=SIFT score for rsID is predicted to be deleterious by the Variant Effect Predictor tool (Ensembl), PROVEAN-D: YES=PROVEAN score is predicted to be deleterious by the Variant Effect Predictor tool (Ensembl), PolyPhenHVAR=D: YES=PolyPhenHVAR score is predicted to be deleterious by the Variant Effect Predictor tool (Ensembl), Clinical_Significance (CLINVAR): clinical significance for rsID (CLINVAR/ Variant Effect Predictor tool), Trait (CLINVAR): clinical trait correlating with rsID (CLINVAR/ Variant Effect Predictor tool), HS: heterozygosity of the subpopulation, HT: heterozygosity of the total population, FST: fixation index.

**Supplemental Table 1B** – Peptide Ancestry Informative Markers – Functional enrichment analyses (metascape.org).

**Supplemental Table 2** – pAIMs Quantified and genotype for rsID encoding pAIM - Endometrial cancer cell Lines (n=13). UniprotAccMatch: Alternate protein accessions mapping to pAIMs (Uniprot.org). GT: genotype, GT=1_1 (homozygous for variant encoding pAIM), GT=0_1 (heterozygous for pAIM), GT=0_0 (homozygous for reference).

**Supplemental Table 3** – pAIMs Quantified - Uterine Leiomyomas (Discovery, n=66). UniprotAccMatch: Alternate protein accessions mapping to pAIMs (Uniprot.org).

**Supplemental Table 4** – pAIMs Quantified and genotype for rsID encoding pAIM - Uterine Leiomyomas (Discovery, n=26). UniprotAccMatch: Alternate protein accessions mapping to pAIMs (Uniprot.org). GT: genotype, GT=1_1 (homozygous for variant encoding pAIM), GT=0_1 (heterozygous for pAIM), GT=0_0 (homozygous for reference).

**Supplemental Table 5 –** Parallel reaction monitoring assay of SERPINA1-V237A and CPPED1-A19D pAIM variants. Table details the dot product similarity (dotp) as well as sum intensities for the top five y-ions quantified by targeted, tandem mass spectrometry for endogenous as well as stable isotope standard (Heavy SIS) peptides in triplicate, technical replicate injections of fibroid FFPE tissue digests from African-American (n=3) and European-American (n=3) women.

## Notes

Funding Statement: This study was supported in part by the U.S. Department of Defense - Uniformed Services University of the Health Sciences (HU0001-16-2-0006 and HU0001-16-2-0014)

